# *Fusarium culmorum* produces NX-2 toxin simultaneously with deoxynivalenol and 3-acetyl-deoxynivalenol or nivalenol (submitted to Toxins)

**DOI:** 10.1101/2022.06.10.495629

**Authors:** Simon Schiwek, Mohammad Alhussein, Charlotte Rodemann, Tuvshinjargal Budragchaa, Lukas Beule, Andreas von Tiedemann, Petr Karlovsky

## Abstract

*Fusarium culmorum* is a major pathogen of grain crops. Infected plants accumulate deoxynivalenol (DON), 3-acetyl-deoxynivalenol (3-ADON), or nivalenol (NIV), which are mycotoxins of the trichothecene B group. These toxins are also produced by *F. graminearum* species complex. New trichothecenes structurally similar to trichothecenes B but lacking the carbonyl group on C-8, designated NX toxins, were recently discovered in atypical isolates of *F. graminearum* from North America. Only these isolates and a few strains of a yet to be characterized *Fusarium* species from South Africa are known to produce NX-2 and other NX toxins. Here we report that among 20 *F. culmorum* strains isolated from maize, wheat, and oat in Europe and Asia over a period of 70 years, 18 strains produced NX-2 simultaneously with 3-ADON and DON or NIV. Rice cultures of strains producing 3-ADON accumulated NX-2 in amounts corresponding to 2-8% of 3-ADON (1.2 - 36 mg/kg). A strain producing NIV accumulated NX-2 and NIV at comparable amounts (13.6 and 10.3 mg/kg, respectively). In *F. graminearum*, producers of NX-2 possess a special variant of cytochrome P450 monooxygenase encoded by *TRI1* that is unable to oxidize C-8. In *F. culmorum*, producers and non-producers of NX-2 possess identical *TRI1*; the reason for the production of NX-2 is unknown. Our results indicate that production of NX-2 simultaneously with trichothecenes B is a common feature of *F. culmorum*.

**Key result:** Isolates of *Fusarium culmorum* obtained from different hosts at different locations over several decades produced NX-2, which is a type A trichothecene structurally related to 3-ADON.

## 1. Introduction

*Fusarium* head blight (FHB) is a cosmopolitan disease of small-grain cereals with a high economic impact [1] caused by several Fusarium species. *F. graminearum* and *F. culmorum* belong to major causal agents of FHB [2]. Infection of grain crops with *Fusarium* spp. causes yield losses and contamination of grains with toxic metabolites (mycotoxins), impairing food safety [3-4].

*F. culmorum* belongs to the *F. sambucinum* species complex [5]. It infects a wide range of grain crops such as wheat [2,6], maize [7-9], barley [10,11], oat [10], triticale [11], and rye [11]. *F. culmorum* typically co-occur with other *Fusarium* species. In most reports, the incidence of *F. culmorum* in crops was second to *F. graminearum*, but in some crops, growing regions, and years, *F. culmorum* dominated [2,6,10]. Infected plant material often contains trichothecene mycotoxins deoxynivalenol (DON), nivalenol (NIV), their acetylated derivatives 3-acetyl-deoxynivalenol (3-ADON) and 15-ac-etyl-deoxynivalenol (15-ADON), fusarenon X (4-acetyl-nivalenol), and zearalenone (ZEN) [4,12].

Biosynthesis of trichothecenes in *Fusarium* spp. is encoded by *TRI* genes, comprising the *TRI1*-*TRI1*6 cluster and the *TRI101* gene located outside of the cluster [13]. According to the dominant trichothecene produced, *Fusarium* strains producing trichothecenes B are partitioned into the 3-ADON chemotype, the 15-ADON chemotype, and the NIV chemotype [13].

The distribution of chemotypes among wheat-producing regions exhibits a strong geographic pattern with most areas dominated by either 3-ADON or 15-ADON chemotype (reviewed in [15]; a succinct overview can also be found in the Introduction of [16]). The 3-ADON chemotype in the USA was assumed to be introduced from Europe [17]. Some studies have not found any relationship between chemotype and aggressiveness of *F. graminearum* [18,19], but other studies found the 3-ADON chemotype more aggressive than the 15-ADON chemotype [20-23]. As several authors suggested [20,21], the discrepancy between earlier and later studies may be accounted for by the use of different inoculation methods because DON is not required for initial infection but it facilitates spread of the pathogen along the spike. Greater aggressiveness of the 3-ADON chemotype as compared to the 15-ADON chemotype is in line with a shift of *F. graminearum* populations from 15-DON to 3-ADON producers in North America over the last two decades [16,21]. In several studies of the population structure of *F. graminearum*, fitness advantage of 3-ADON producers due to their higher aggressiveness was postulated and correlations with fitness parameters were determined. The reason for a greater aggressiveness of 3-ADON producers as compared to 15-ADON producers has rarely been addressed. A plausible hypothesis was coined by Poppenberger et al. [24] based on their finding that 3-ADON was protected against glucosylation by UDP-glucosyltransferase from *Arabidopsis thaliana* (see also Section 3.5.).

Trichothecene chemotype has been monitored extensively in *F. graminearum* (e.g., [9,16,18-23,25]; reviewed in [15]). Fewer studies monitored the chemotype in *F. culmorum*, which only comprises 3-ADON and NIV types [9,26-28]. In the past, chemotypes were often assigned according polymorphisms in *TRI* genes [25], but frequently reported discrepancies between the chemotype prediction by PCR-RFLP and results of chemical analysis have questioned this approach [14,16,29].

During an investigation of *F. graminearum* strains colonizing wheat heads in North America, a new type A trichothecene mycotoxin, the NX-2, was discovered [30]. The structure of NX-2 is similar to 3-ADON but it lacks a carbonyl group at C-8. Deacetylated derivative of NX-2, called NX-3, has also been reported. Very recently, these metabolites have also been found in a yet to be characterized species of the *Fusarium sambucinum* species complex from South Africa [31]. The first strains of *F. graminearum* from North America producing NX toxins did not produce any trichothecene type B under laboratory conditions. The lack of known trichothecenes actually motivated the investigation that led to the discovery of NX toxins [30]. Strains producing NX-2 were originally found at very low frequency but a recent study by Lofgren et al. [32] estimated that 20% of the *F. graminearum* population in the USA produce NX-2. An even more recent study of *F. graminearum* strains in Canada [16] identified producers of NX-2 (which they designated 3ANX for consistency with the nomenclature of ADONs) among strains of *F. graminearum* assigned to the 15-ADON chemotype. Contrary to the reports that NX-2-producering strains from the USA have not produced DON, NIV, or their acetylated derivatives [30,33], most strains in their work [16] produced NX-2 simultaneously with 15-ADON.

Here we report that most isolates of *F. culmorum* collected in Europe and Asia produced NX-2 toxin simultaneously with DON, 3-ADON, or NIV.

## 2. Results

### 2.1. Mycotoxin production in rice cultures

Our set of 20 *F. culmorum* strains consisted of 14 strains isolated from maize, wheat, and oat in Germany and 6 strains obtained from other laboratories and culture collections, which were isolated in different countries in a time span of 70 years. The analysis of rice culture extracts revealed accumulation of NX-2 at concentrations larger than 1 mg/kg in 18 cultures (**Table 1**). All strains except one were primarily 3-ADON producers; one isolate was assigned to the NIV chemotype. The amounts of NX-2 and 3-ADON in cultures of strains of the 3-ADON chemotype were tightly correlated (**Figure 1**). NX-2 in these cultures accumulated to levels corresponding to 2-8% of 3-ADON. In the culture of the only isolate of the NIV chemotype (isolate 59.6st), the concentrations of NIV and NX-2 were comparable (**Table 1**).

**Table 1.** Accumulation of trichothecenes in rice cultures of *F. culmorum* strains.

| Isolate | Mycotoxin content (mg/kg) |  |  |  |  |
| --- | --- | --- | --- | --- | --- |
|  | NIV | DON | 15-ADON | 3-ADON | NX-2 |
| 31.6st | 0.04 | 25.6 | <LOD | 156 | 3.2 |
| 215.1st | <LOD | 4.2 | <LOD | 90.4 | 2.4 |
| 227.2cst | <LOD | 0.5 | <LOD | 2.1 | <LOD |
| 240.2sp | 0.38 | 355 | <LOD | 552 | 35.6 |
| IPP0211 | 0.31 | 566 | <LOD | 412 | 12.2 |
| IPP0212 | 0.10 | 162 | <LOD | 463 | 16.4 |
| IPP0213 | 0.33 | 639 | <LOD | 388 | 9.4 |
| IPP0618 | <LOD | 20.3 | <LOD | 175 | 3.7 |
| IPP0619 | <LOD | 15.9 | <LOD | 272 | 14.4 |
| IPP0999 | <LOQ | <LOD | <LOD | <LOD | <LOD |
| IPP1000 | <LOD | 12.6 | <LOD | 132 | 2.7 |
| DSM62184 | 0.05 | 108 | <LOD | 173 | 13.5 |
| 966 | <LOD | 25.7 | <LOD | 118 | 3.2 |
| 969 | <LOQ | 14.3 | <LOD | 147 | 3.2 |
| 3.37 | <LOD | 9.0 | <LOD | 68.6 | 1.4 |
| DSM62188 | <LOQ | 46.5 | <LOD | 294 | 13.6 |
| 55.6st | <LOQ | 31.3 | <LOD | 85.9 | 1.2 |
| 59.6st | 13.6 | 0.4 | 0.9 | 0.63 | 10.3 |
| K11.2 | <LOD | 1.0 | <LOD | 46.3 | 1.2 |
| J31.2 | 0.08 | 127 | <LOD | 447 | 18.6 |

LOD = limit of determination; LOQ = limit of quantification.

**Figure 1.**
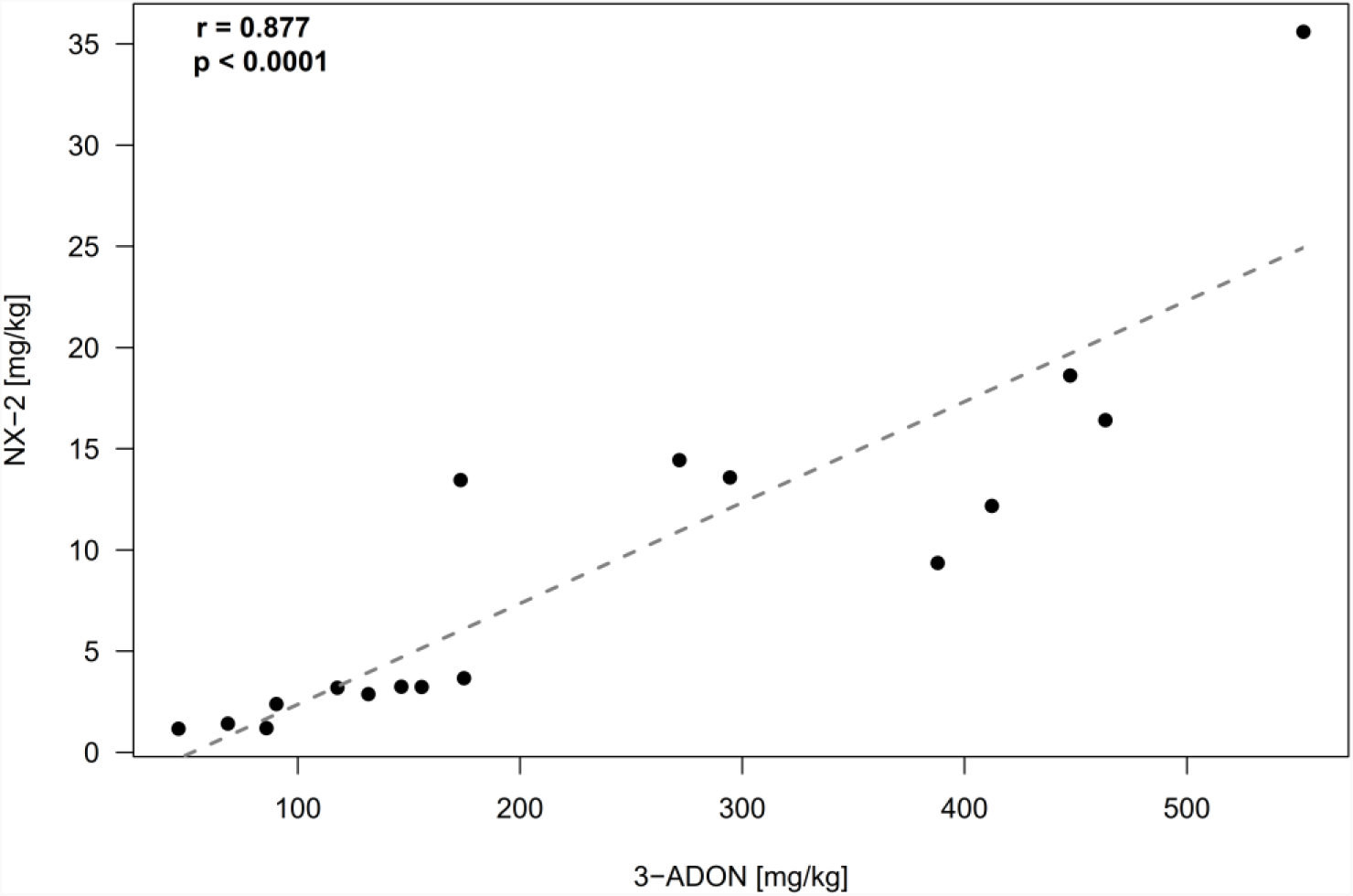
Relationship (Pearson correlation) between NX-2 and 3-ADON concentrations in rice cultures of *F. culmorum*. Each culture (**Table 1**) is represented by a single data point (black dots). Isolate 59.6st, which was assigned to the NIV chemotype, and isolates 227.2 and IPP0999, in which the concentrations of NX-2 were below the limit of detection, were excluded.

### 2.2 Confirmation of the structure of NX-2

Putative NX-2 accumulating in rice cultures of *F. culmorum* was originally identified by comparing its retention time in HPLC and MS/MS fragmentation with data obtained with purified NX-2 from Prof. Franz Berthiller (BOKU Vienna, Austria). Because production of NX-2 by *F. culmorum* has not been reported before and an isomer of NX-2 could possess the same retention time and generate product ions with the same m/z values, we purified putative NX-2 from rice cultures of *F. culmorum* 240.2sp (**Figure 2**) to verify its structure. Approx. 5 mg of pure metabolite were obtained from 480 g of dry rice culture (see Section 5.4 for details).

**Figure 2.**
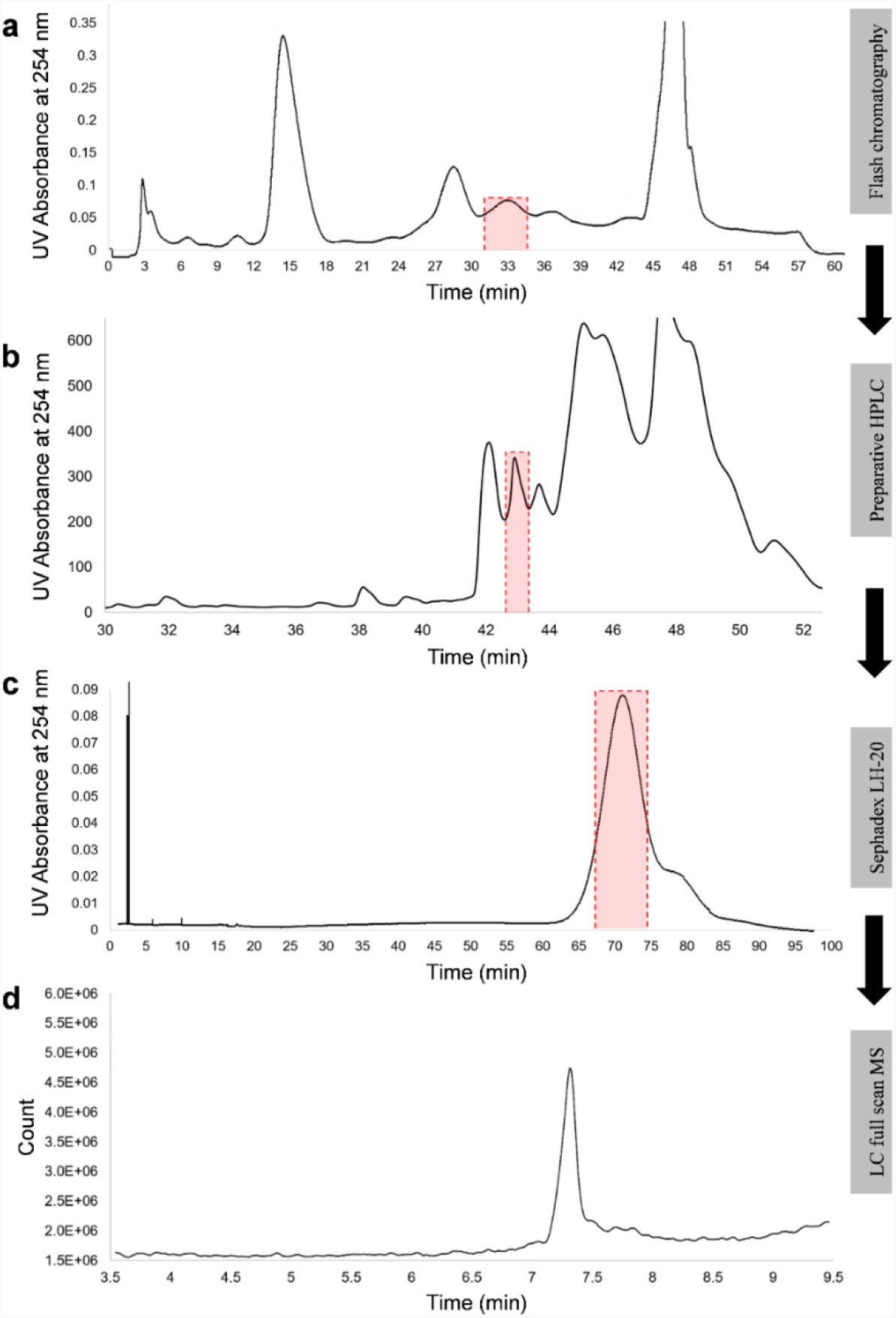
Purification of NX-2 toxin from rice culture of *F. culmorum* 240.2sp. Three weeks-old rice cultures were extracted with methanol/water/acetic acid and putative NX-2 was enriched by chromatography on C18 cartridge (a), polar-modified C18 column (b), and Sephadex LH-20 (c). Purity of the metabolite was established by HPLC-MS in a full-scan mode (d).

One-(^1^H and ^13^C) and two-dimensional (^1^H,^13^C-HSQC, ^1^H,^13^C-HMBC, ^1^H,^1^H-COSY) NMR spectroscopic analysis was performed on the purified metabolite (**Supplementary Figure S1-S6**). The spectroscopic data were in accordance with published data for NX-2 (**Supplementary Table S1**). The structure of NX-2 is shown in **Figure 3**.

**Figure 3.**
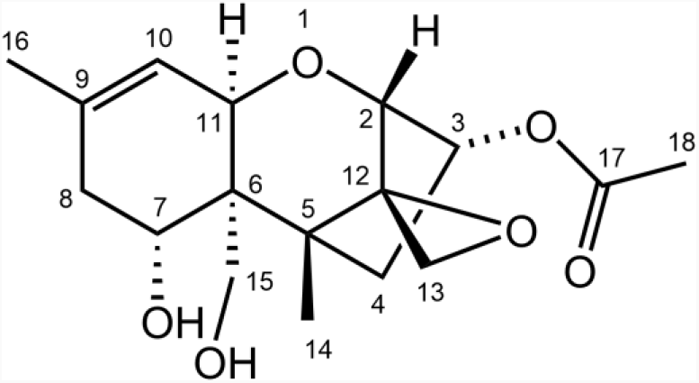
Structure of NX-2 toxin.

### 2.3. Species assignment and investigation of polymorphisms in TRI1

Examination of spores assigned all isolates analyzed for trichothecene production (**Table 1**) to *Fusarium culmorum*. Morphological characterization was complemented by the analysis of melting curves of amplicons of taxonomically informative genes encoding translation elongation factor 1α (TEF*-1α*) and the second largest component of the RNA polymerases II (*RPB2*) [34]. Species identification was further strengthened by the analysis of full-length sequence (1,753 nt) of the *TRI1* gene, which encodes cytochrome P450 monooxygenase catalyzing oxygenation of calonectrin on C-7 and C-8. The sequences of *TRI1* obtained from isolates of *F. culmorum* used in this study (accession numbers OM144918 to OM144937) were aligned with a set of reference sequences (**Supplementary Table S3**) and used for phylogenetic analysis by the maximum-likelihood method. Separation of *F. culmorum* from other *Fusarium* species was highly supported, indicating that *TRI1* is taxonomically informative in *Fusarium* species producing trichothecenes.

The reason for selecting the *TRI1* gene for the analysis was that the product of *TRI1* catalyzes biosynthetic steps distinguishing NX toxins from trichothecenes B, and that polymorphisms in *TRI1* differentiating *F. graminearum* strains producing NX toxins from nonproducers were identified [30]. As expected, all *F. graminearum* strains not producing NX toxins were separated from all strains producing NX toxins. In contrast, no polymorphisms separating *F. culmorum* strains producing NX toxins from nonproducers were found in the *TRI1* gene (**Figure 4**).

**Figure 4.**
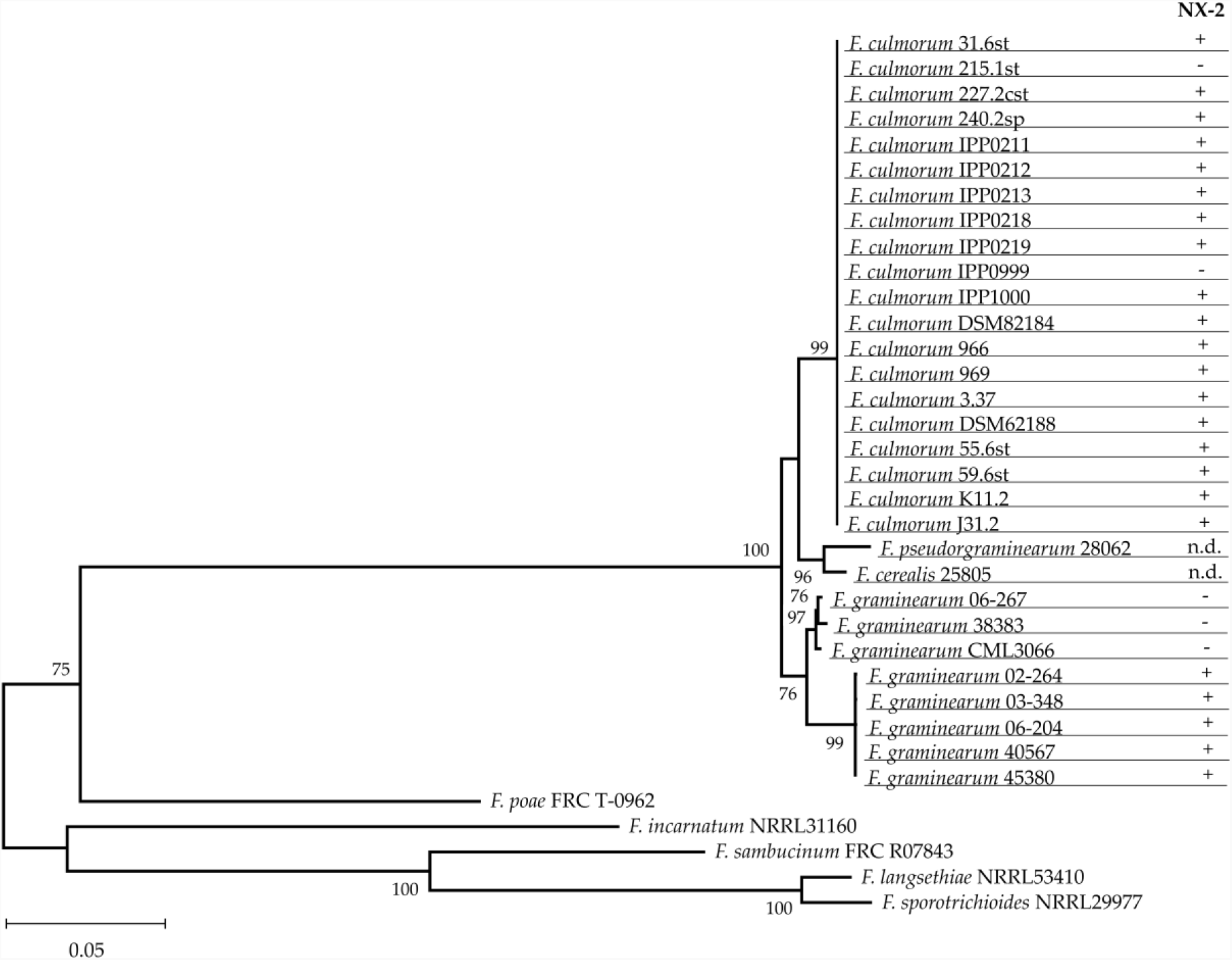
Maximum likelihood estimate of phylogenetic relationships among *TRI1* genes in *Fusarium* spp. Complete sequences of *TRI1* genes of the investigated isolates of *F. culmorum* (see **Table 3**) and reference sequences were subjected to maximum likelihood analysis [36] assuming Tamura-Nei model [37]. Bootstrap values (1,000 replications) are shown next to the nodes. Production of NX-2 was determined by HPLC-MS/MS; n.d. stands for no data available. Nucleotide sequences were deposited at NCBI with accession nos. listed in **Supplementary Table S3**.

Amino acid sequences of translation products of *TRI1* genes, designated Tri1, were identical for all isolates of *F. culmorum* in this study. Isolates of *F. graminearum* producing NX toxins differed from nonproducers in 14 amino acid residues within the heme-binding motif [35] (**Table 2**). In all *F. culmorum* strains used in this study, comprising 18 producers and 2 nonproducers of NX-2, these amino acid residues were identical, and they matched the corresponded residues in strains of *F. graminearum* that did not produce NX toxins. Thus, the reason for the production of NX-2 by *F. culmorum* is not an NX-specific form of *TRI1* found in NX-2 producers of *F. graminearum*.

**Table 2.**
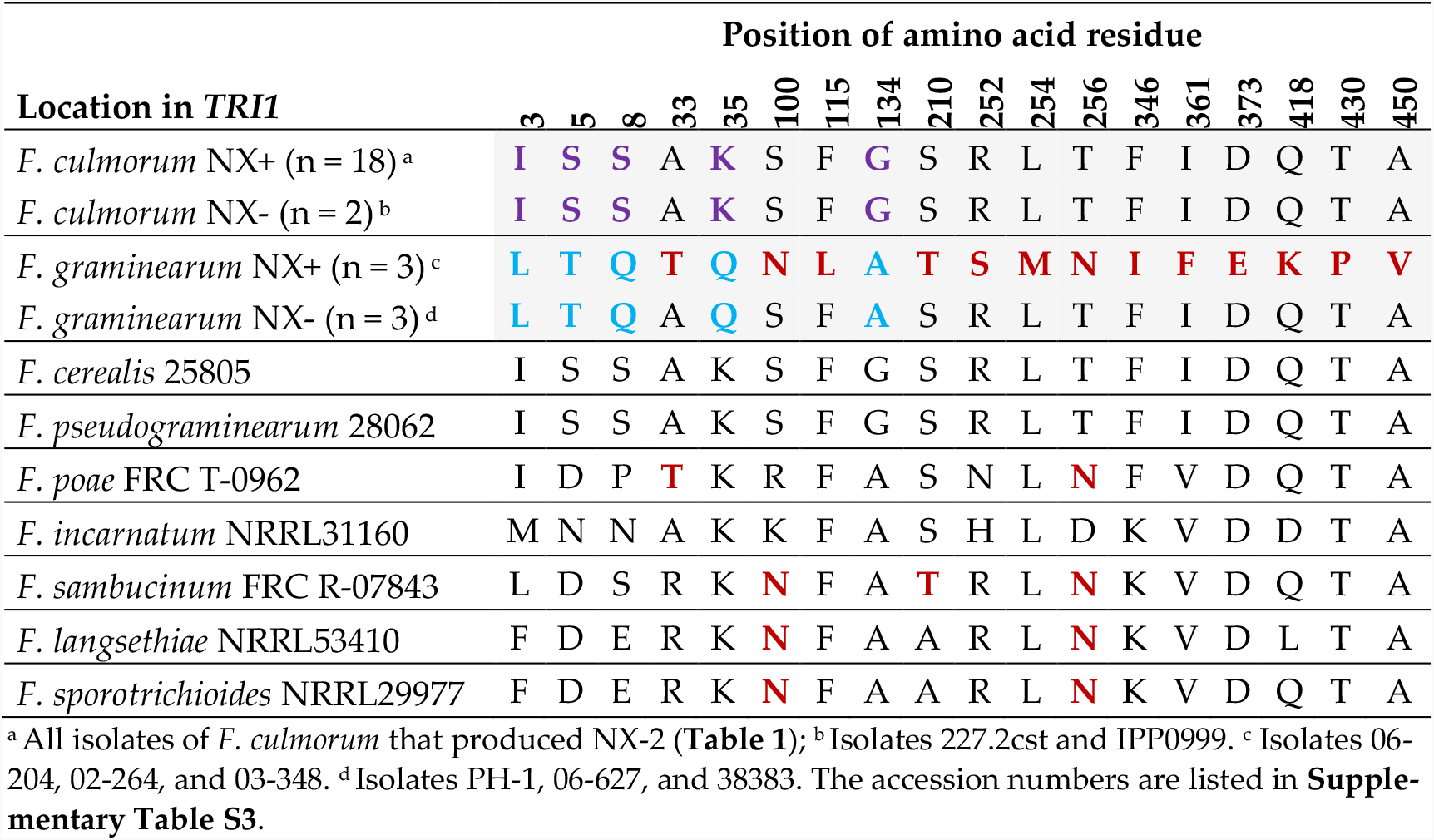
Amino acid residues specific for the production of NX-2 in the translation product of *TRI1* of *F. graminearum* and corresponding residues in *F. culmorum*. Species-specific positions are highlighted in purple for *F. culmorum* and blue for *F. graminearum*. Positions reported to distinguish NX-2-producing strains of *F. graminearum* [35] are marked red.

**Table 3.** Strains of *F. culmorum* used in this study.

| Species | Origin | Year | Isolate | Host |
| --- | --- | --- | --- | --- |
| *F. culmorum* | Germany | 2017 | 31.6st | Maize stalk |
| *F. culmorum* | Germany | 2018 | 215.1st | Maize stalk |
| *F. culmorum* | Germany | 2018 | 227.2cst | Maize stalk |
| *F. culmorum* | Germany | 2018 | 240.2sp | Maize rachis |
| *F. culmorum* | - | < 1984<sup>a</sup> | IPP0211<sup>b</sup> | Wheat |
| *F. culmorum* | Italy | < 1993<sup>a</sup> | IPP0212<sup>b</sup> | Wheat |
| *F. culmorum* | - | 1993 | IPP0213<sup>b</sup> | Barley |
| *F. culmorum* | Hungary | 1991 | IPP0618<sup>b</sup> | Wheat |
| *F. culmorum* | Germany | 1991 | IPP0619<sup>b</sup> | Wheat |
| *F. culmorum* | Germany | 2010 | IPP0999<sup>b</sup> | Wheat |
| *F. culmorum* | Germany | 2010 | IPP1000<sup>b</sup> | Wheat |
| *F. culmorum* | Germany | 1952 | DSM62184<sup>c</sup> | Maize grain |
| *F. culmorum* | Syria | 2009-2010 | 966<sup>d</sup> | Wheat |
| *F. culmorum* | Syria | 2009-2010 | 969<sup>d</sup> | Wheat |
| *F. culmorum* | Germany | 2004 | 3.37<sup>b</sup> | Wheat |
| *F. culmorum* | Germany | 1990 | DSM62188<sup>c</sup> | Maize stalk |
| *F. culmorum* | Germany | 2017 | 55.6st | Maize stalk |
| *F. culmorum* | Germany | 2017 | 59.6st | Maize stalk |
| *F. culmorum* | Germany | 2021 | K11.2 | Oat |
| *F. culmorum* | Germany | 2021 | J31.2 | Oat |

<sup>a</sup> Isolates were collected before the specified year. <sup>b</sup> Isolates of the Plant Phytopathology and Crop Protection Section at the University of Göttingen, Göttingen, Germany. <sup>c</sup> Strains were obtained from the German Collection of Microorganisms and Cell Cultures, Braunschweig, Germany.
<sup>d</sup> The isolates were described by Alkadri et al. [40].

## 3. Discussion

### 3.1. Production of NX-2 is a characteristic feature of *F. culmorum*

So far, production of NX-2 has exclusively be reported in *F. graminearum*, which shares a high level of genomic similarity with *F. culmorum*, produces the same mycotoxins, and has the same host range [6,38]. The production of NX-2 was discovered in an atypical population of *F. graminearum* from the Midwest of the U.S.A., named Northland population, which did not produce any known trichothecene [33]. Producers of NX-2 were later found in this population [30] but only a small fraction of *F. graminearum* strains collected in the area produced NX toxins. A later study on *F. graminearum* in wheat in Ontario (Canada), however, reported that 80% of investigated strains produced NX toxins [16]. A key finding of the current study is that NX-2 production is not limited to *F. graminearum*, and, contrary to the previous assumption [35], it is not endemic to Northern USA and Southern Canada.

The ability to produce NX-2 appears to be a common feature of *F. culmorum*. Among 20 strains of *F. culmorum* isolated in Europe and Asia over a period of 70 years, 18 strains produced NX-2 (**Table 1**). We suppose that the production of NX-2 toxins in cultures of *F. culmorum* remained unnoticed because NX toxins were not monitored in routine surveys. No commercial standards for NX-2 are available at the time of writing. Because infection of grain crops with *F. culmorum* is widespread, we assume that contamination of grains and grain products with NX-2 might be common.

### 3.2. Both chemotypes of *F. culmorum* produce NX-2

Isolates of *F. culmorum* belong to the 3-ADON and NIV chemotypes; 15-ADON chemotype is absent [26-28,39,40]. Most *F. culmorum* isolates studied in this work belonged to the 3-ADON chemotype and produced NX-2 toxin. The only strain of the NIV chemotype produced NX-2 toxin, too (**Table 1**). Investigation of further isolates has yet to confirm that this generally holds for strains of the NIV chemotype. In *F. graminearum*, all producers of NX toxins identified in the first report from the USA belonged to the 3-ADON chemotype [30]. Another group studying *F. graminearum* strains from Canada confirmed this finding: NX-2 producers were rare, and all of them (including 5 isolates from Ontario, see below) were assigned to the 3-ADON chemotype [41]. The most recent study on *F. graminearum* from Ontario reported contradictory results, assigning most NX-2 producers to the 15-ADON chemotype [16]. Production of NX-2 was verified by chemical analysis in both studies. In the study of Kelly et al. [41], the 3-ADON/15-ADON chemotype was assessed by PCR-RFLP, while Crippin at al. [16] used both PCR-RFLP and chemical analysis. Incongruencies between the chemotype prediction by PCR-RFLP and the results of chemical analysis, reported by several studies [14,16,29,42], cannot explain the contradictory chemotype assignments mentioned above because both studies used the same PCR assay (*TRI*3/*TRI*12), yet Crippin et al. [16] reported that all their NX-2 producers belonged to the 15-ADON chemotype while Kelly at al. [41] assigned all their NX-2 producers to the 3-ADON chemotype. Crippin at al. [16] also analyzed trichothecenes in cultures on three growth media by HPLC-MS/MS, using a special elution gradient for the separation of 3-ADON and 15-ADON. (These mycotoxins often co-elute, and they cannot be reliable distinguished by MS fragmentation). They reported the concentrations of 15-ADON, but unfortunately not the concentrations of 3-ADON and/or DON.

The acetylation of C-3-OH of NX-2 (**Figure 3**) is reminiscent of 3-ADON. Varga at al. [30] showed that when part of the translation product of *TRI1* in a strain of the 15-ADON chemotype was replaced with a *TRI1* segment specific for the production of NX-2, the recombinant strain accumulated NX-4 (a derivative acetylated on C-15-OH, reminiscent of 15-ADON) rather than NX-2. The replacement of NX-specific polypeptide in an NX-2 producer with a protein segment from a 15-ADON-producer converted the NX-2-producing strain into a 3-ADON producer. Both results strongly support the hypothesis that NX-2 producers in *F. graminearum* developed from strains of the 3-ADON chemotype. The controversy between the assignment of chemotypes to NX-2 producers from Ontario in [41] and [16] remains unresolved.

### *3*.*3. F. culmorum* produces NX-2 simultaneously with DON and 3-ADON or NIV

A search for new trichothecenes in *F. graminearum* that facilitated the discovery of NX toxins was motivated by a failure to detect any trichothecene in a group of atypical strains of *F. graminearum* from the USA [30,33]. These strains produced NX toxins but no trichothecenes B. The recent study from Ontario cited above [16] reported that cultures of *F. graminearum* accumulating NX-2 simultaneously accumulated 15-ADON.

Cultures of NX-2-producing *F. culmorum* strains in our study accumulated 3-ADON (most strains) or NIV (a single strain). In cultures of the 3-ADON chemotype, NX-2 accumulated at amounts corresponding to 2-8% of 3-ADON, and the content of NX-2 and trichothecenes B tightly correlated (**Figure 1**). In contrast, the concentrations of NX-2 to 15-ADON in *F. graminearum* cultures in the study of Crippin et al. [16] were not correlated; the ratio NX-2/15-ADON varied from 0.003 to 33.

The only strain in our work producing NX-2 at an amount comparable to trichothecenes B was a strain of the NIV chemotype (**Table 1**). In rice cultures of this strain, NX-2 reached 75% of the concentration of NIV and 66% of total trichothecenes B. Future studies have to clarify whether relatively high production of NX-2 is a typical feature of strains of *F. culmorum* with the NIV chemotype. In *F. graminearum*, no strain of the NIV chemotype have so far been reported to produce NX-2.

### 3.4. NX-2 production by *F. culmorum* is not caused by a variant of TRI1

The reason for the production of NX toxins by certain strains of *F. graminearum* is that these strains harbor a special variant of the *TRI1* gene [30]. The gene, which is only present in genomes of trichothecene-producing *Fusarium* species [43], encodes cytochrome P450 monooxygenase (calonectrin C7/C8 hydroxylase) Tri1 that catalyzes oxidation of C-7 and C-8 [44,45]. The difference between *TRI1* of NX-2 producers and nonproducers allowed a PCR-RFLP assay for NX-2 producers [46]. We have not found such a polymorphism in the *TRI1* gene of *F. culmorum*.

In *F. graminearum*, changes in Tri1 specific for NX toxins occurred in the heme binding motif close to the C-terminus. Furthermore, Ramdass et al. [47] found an new potential glycosylation site in the enzyme of NX-2 producers. It was located at a large distance from the heme binding site but the authors suggested that glycosylation may modulate enzyme activity via protein folding. None of these changes occurred in the *TRI1* gene of NX-2-producing *F. culmorum*. Some residues in the heme-binding segment differed from corresponding residues in *F. graminearum* (purple in **Table 2**), but these residues occurred in NX-2 producers as well as nonproducers.

Factors other than the amino acid sequence of Tri1 must suppress its activity towards C-8 of calonectrin in *F. culmorum* strains producing NX-2. The enzyme is located inside endoplasmic reticulum with two hydrophobic segments close to the ends of the protein crossing the membrane [47]. Other proteins and especially other cytochromes P450, which compete with Tri1 for the same NADPH-cytochrome P450 reductase, share this location. Cytochromes P450 are known to interact with each other and form heterooligomers, which modifies their activity [48]. Interaction of Tri1 with other proteins anchored in the membrane of endoplasmic reticulum may suppresses the oxidation of C-8 in NX-2-producing strains of *F. culmorum*.

### 3.5. NX-2 and the aggressivenss of *F. graminearum* and *F. culmorum*

In *F. graminearum*, a shift from 15-ADON to 3-ADON chemotype observed in the last decades supports the hypothesis that the 3-ADON chemotype is more aggressive. For instance, an increase of the 3-ADON chemotype at locations where 15-ADON producers used to be predominant was reported in a study from North America [49]. Many studies of the population structure in *F. graminearum* elucidated the relationship between genotype, chemotype and aggressiveness, aiming to explain how selection and gene flow shaped the populations (e.g., 17,20,35,41,46,49,51,53]. Field studies documented the success of the 3-ADON chemotype of *F. graminearum* but they could not address its cause, which requires a biochemical approach. According to a hypothesis from the lab of Gerhard Adam [24], acetylation of C-3-OH prevents detoxification of DON, which is a virulence factor, by plant UDP-glycosyltransferases. In NX-2, the hydroxyl on C-3 is also protected by acetylation, and Varga at al. [30] suggested that the production of NX-2 may benefit *F. graminearum* during the infection in the same way. Glucosylation of DON takes place in the cytoplasm while the target of DON is protein synthesis in rough endoplasmic reticulum, where the acetyl group would have to be removed. This is plausible because endoplasmic reticulum is rich in hydrolases [50]. Varga et al. [15] also speculated that the lack of carbonyl on C-8 circumvents detoxification by glutathionylation [15]. This hypothesis holds for *F. cumorum*, too.

Can the effect of NX-2 on the aggressiveness of *F. graminearum* or *F. culmorum* be proved? Field trials with natural isolates are unlikely to generate a conclusive proof. The same situation exists for the claim that the 3-ADON chemotype is more aggressive than the 15-ADON chemotype, which was supported by some studies [20-23] yet rejected by others [18,19,53]. Field isolates differ in many properties modulating aggressiveness. Some of them are likely linked to the chemotype. Isogenic strains differing only in the gene in question are required. Regarding relative aggressiveness of the chemotypes 3-ADON and 15-ADON, strains with swapped *TRI8* genes or *TRI8* chimeras, constructed by Alexander et al. [57], could be used. Similarly, infection experiments with *F. graminearum* strains harboring *TRI1* with swapped domains controlling the NX-2 production, constructed by Varga at al. [30], would reveal the effect of NX-2 production on aggressiveness.

### 3.6. Do further *Fusarium* species produce NX toxins?

Comparison of the amino acid residues in Tri1 sequence distinguishing NX-2-producing isolates from nonproducers in *F. graminearum* (**Table 2**) showed that certain residues specific for the production of NX-2 are also present in Tri1 of other *Fusarium* species. For instance, Tri1 in a particular *F. sambucinum* strain shared 3 residues with NX-2 producing strains of *F. graminearum*, and it differed from Tri1 of strains not producing NX-2 in other 3 positions. We suggest that this species be examined for NX-2 production. *F. cerealis* and *F. pseudograminearum* harbor *TRI1* genes very similar to *TRI1* of *F. culmorum* (**Figure 4**), and the amino acid sequence in the heme-binding segment of their Tri1 protein is identical with the corresponding sequence in *F. culmorum* (**Table 2**). Examination of these species for the production of NX toxins, too, appears worthwhile.

### 3.7. Can a single strain of *F. culmorum* produce both DON and NIV?

It is generally assumed that *F. culmorum* and *F. graminearum* produces either NIV or DON but not both. We believe that this is view is biased by low sensitivity of analytical methods in the past, which only detected trichothecenes that were present at high concentrations (e.g., [27]). Accounts of the production of both trichothecenes by a single strain have been largely overlooked. For instance, Foroud et al. [3] write in their recent review “ NIV chemotypes do not produce DON”, though many studies, such as [58,20,59] (the first two of which are even cited in [3]), reported simultaneous production of DON and NIV by single strains of *F. culmorum* or *F. graminearum*.

NIV and DON have a common precursor. In NIV producers, the precursor is hydroxylated at C-4 by Tri13 [57]. We speculate that before the entire pool is hydroxylated, some enters the path leading to DON. Therefore, we assume that all cultures producing NIV contain small amounts of DON. The presence of NIV in cultures of DON producers can be explained by a residual activity of Tri13 or by the presence of hydroxylases with a relaxed substrate specificity. In line with this reasoning, cultures of NIV-producing strain 59.6st in our work accumulated relatively large amounts of DON while cultures of most DON producers contained small amounts of NIV (Table 1). We expect that with the widespread use of sensitive analytical methods, small amounts of NIV will be often found in cultures of *Fusarium* strains producing DON, and substantial amounts of DON will be found in cultures of all strains producing NIV.

## 4. Conclusions

Most strains of *Fusarium culmorum* produce NX-2 toxin simultaneously with deoxynivalenol, 3acetyldeoxynivalenol, or nivalenol. The strains producing NX-2 do not possess a specific variant of the *TRI1* gene known from NX-2-producing strains of *F. graminearum*.

## 5. Materials and Methods

### 5.1. Fungal strains

Strains of *F. culmorum* are listed in **Table 3**. Strains isolated in the course of this study were obtained from maize grains, rachis, stalks, and from oat grains according to Leslie and Summerell [54]. Briefly, samples were surface sterilized for 10 min with 0.1% silver nitrate or 3% sodium hypochlorite, rinsed, and placed on potato dextrose agar (PDA). Isolates were purified via single-spore cultures and grown on PDA for colony characteristics and on low-nutrient agar (SNA, [55]) under long-wave UV light for morphological characterization of spores. For long-term storage, fungal cultures were freeze-dried.

### 5.2 DNA methods

Fungal DNA was extracted using a CTAB-based protocol [56] from 10 mg of lyophilized mycelium and dissolved in 50 µl TE (10 mM Tris, 1 mM EDTA, pH 8.0). Segments of marker genes *TEF1α* and *RPB2* were amplified and the PCR products subjected to high-resolution melting curve (HRM) analysis as described previously [34]. The *TRI1* gene was amplified for sequencing as four overlapping fragments (**Table 4**), which were sequenced by the Sanger method and assembled to full-length gene sequences. PCR was carried out in a peqSTAR 96 thermocycler (PEQLAB, Erlangen, Germany) in a total reaction volume of 25 µl. Reaction mixtures were composed of a buffer (10 mM Tris-HCl, 50 mM KCl, 1.5 mM MgCl_2_, pH 8.3 at 25°C; New England Biolabs, Beverly, MA, USA) adjusted to 2 mM final MgCl_2_ concentration containing 100 µM of each deoxyribonucleoside triphosphate (Bioline, Luckenwalde, Germany), 0.3 µM of each primer, 0.62 U HotStart-polymerase (New England Biolabs, Beverly, Massachusetts, USA), and 1 µL template DNA solution diluted 100-times. The thermocycling conditions are specified in **Supplementary Table S2**. PCR products were precipitated with isopropanol, washed with 80% ethanol and sequenced at the facilities of Macrogen Europe (Macrogen Europe, Amsterdam, The Netherlands). The sequences were quality-trimmed with Chromas version 2.6.6 (Technelysium Pty. Ltd., South Brisbane, Australia) and assembled to fulllength gene sequences; the accession numbers are listed in **Supplementary Table S3**. Multiple sequence alignments were performed with ClustalW [60] in MEGAX version 10.1.8 [36]. Phylogenetic relationships among *TRI1* genes in *Fusarium* spp. were investigated by maximum-likelihood analysis using MEGA X under the assumption of Tamura’s and Nei’s substitution model [37].

**Table 4.**
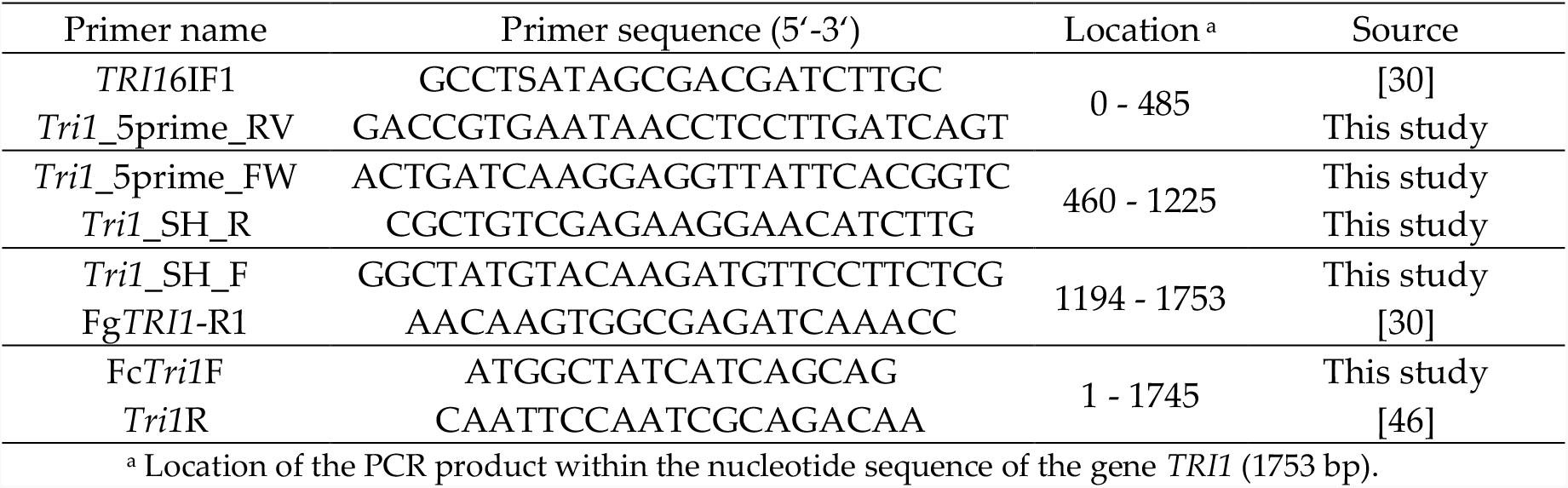
Oligonucleotides used for the sequence analysis of the gene *TRI1*.

### 5.3. Mycotoxin extraction and HPLC-MS/MS

Rice cultures were prepared in 50-ml Falcon tubes (Sarstedt, Nümbrecht, Germany) by autoclaving 3 g dry polished rice with 5 ml tap water. The tubes were inoculated with plugs of PDA (0.5 cm diameter) overgrown with 5-days-old mycelium. Rice medium incubated with agar plugs without mycelium served as a control. The cultures were incubated for 21 days at 25 °C in the dark. Fungal metabolites were extracted by shaking with 30 ml acetonitrile/water/acetic acid (84:15:1 (v/v/v)) overnight. Extracts were dried in a vacuum concentrator (Martin Christ, Osterode am Harz, Germany) and residues re-dissolved in methanol/water (20:80 (v/v)) as described previously [61]. Mycotoxin analysis was carried out using a 1290 Infinity II HPLC system (Agilent Technologies, Waldbronn, Germany) coupled with a 6460 triple quadrupole detector (Agilent Technologies, Waldbronn, Germany). The separation was performed on a Zorbax Eclipse Plus C18 column, 50 × 2.1 mm with 1.8 µm particle size (Agilent Technologies, Waldbronn, Germany). The column oven temperature was 40 °C. Mobile phase A was water with 0.1% formic acid (v/v), and phase B was methanol with 0.1% formic acid (v/v). The gradient was as follows: 0 to 0.2 min, 5% B; 0.2 to 8 min, 5% to 35% B; 8 to 8.5 min, 35% to 98% B; 8.5 to 12 min, 98% B; 12 to 12.5 min, 98% to 5% B; 12.5 to 16 min, 5% B. The calibration curve included 11 concentrations from 0.48 to 500 μg/L. A blank was analyzed after every 7^th^ sample and a quality control standard after every 15^th^ sample. The metabolites were detected in a multiple reaction monitoring (MRM) mode. The acquisition parameters and the limits of detection (LOD) and quantification (LOQ) are listed in **Table 5**.

**Table 5.** Parameters for HPLC-MS/MS analysis of trichothecenes.

| Toxin | Ioniz. mode | Parent ion [m/z] | Fragm. voltage [V] | Collision energy [V] | Product ion [m/z]<sup>c</sup> | LOD<sup>b</sup> [mg/kg] | LOQ<sup>b</sup> [mg/kg] |
| --- | --- | --- | --- | --- | --- | --- | --- |
| NIV | Pos | 313.1 | 93 | 7 | 205.0 | 0.01 | 0.03 |
|  |  |  |  | 15 | <u>175.0</u> |  |  |
| DON | Pos | 297.1 | 100 | 4 | <u>249.1</u> | 0.04 | 0.13 |
|  |  |  |  | 64 | 91.2 |  |  |
| 3-ADON | Pos | 339.2 | 100 | 8 | <u>231.1</u> | 0.02 | 0.08 |
|  |  |  |  | 8 | 203.0 |  |  |
| 15-ADON | Pos | 339.2 | 90 | 10 | 321.1 | 0.07 | 0.23 |
|  |  |  |  | 10 | <u>261.0</u> |  |  |
|  |  |  |  | 10 | 137.0 |  |  |
| NX-2<sup>a</sup> | Pos | 325.2 | 90 | 10 | 247.2 | 0.04 | 0.12 |
|  |  |  |  | 10 | 229.2 |  |  |
|  |  |  |  | 10 | 199.1 |  |  |
|  |  |  |  | 25 | 121.2 |  |  |
|  |  |  |  | 35 | <u>105.1</u> |  |  |

<sup>a</sup> Analytical reference for NX-2 was kindly provided by Dr. Franz Berthiller (BOKU, Vienna, Austria). <sup>b</sup> Limit of detection (LOD) and limit of quantification (LOQ) were calculated based on the standard deviation of the blank [62]. <sup>c</sup> m/z of product ions used for quantification are underlined.

### 5.4. Purification of NX-2

*F. culmorum* isolate 240.2sp, which produced the largest amunts of NX-2, was cultivated on rice media. The cultures were prepared in 4 l flat penicillin flasks with 120 g organic polished rice and 270 ml tap water, which were autoclaved and inoculated with 12 agar-plugs (0.6 cm diameter) from 5-day old fungal cultures. Rice cultures were incubated at 25 °C for three weeks in the dark. NX-2 was extracted with a 5-time excess of methanol/water/acetic acid (90:9:1) by shaking overnight. The extract was concentrated to approximately 15% of its original volume using a rotary evaporator R-100 (Buchi, Flawil, Switzerland). The concentrated extract was partitioned with the same volume of ethyl acetate (EtOAc) three times. Combined EtOAc fractions were dried in a rotary evaporator. The residue was dissolved in methanol/water (30:70) and cleared by centrifugation. The supernatant was subjected to flash chromatography (Sepacore Flash system X10/X50, Buchi, Flawil, Switzerland) equipped with a binary pump (modules C-601/C-605) and an UV detector (module C-635). Metabolites were separated on a reverse-phase column (Chromoband Flash cartridge RP C18 ec 40–63 µm, 262 × 37 mm, Machery-Nagel, Düren, Germany). Mobile phase A was water with 0.2% acetic acid; phase B was methanol with 0.2% acetic acid. The gradient elution program was as follows: 0-2 min, 5% B; 2-42 min, from 5% to 50% B; 42-45 min, from 50% to 98% B; 45-55 min, 98% B; 55-58 min, from 98% to 5% B; 58-66 min, 5% B. The flow rate was 50 ml min^-1^. The separation was monitored at 254 nm. NX-2 fraction was collected at 33 min, the solvent was evaporated, fractions containing NX-2 were re-dissolved in methanol/water (30:70) and subjected to a preparative-HPLC (PU-2086 plus, Jasco, Gross-Umstadt, Germany) equipped with a preparative reverse-phase column (Nucleodur C18 pyramid, 5 µm, 250 × 21mm, Macherey-Nagel, Düren, Germany) and coupled to a UV/VIS detector (UV-970, Jasco, Gross-Umstadt, Germany). NX-2 was eluted by a gradient of water with 0.2% acetic acid (v/v) (solvent A) and methanol with 0.2% acetic acid (solvent B) as follows: 0-5 min, 5% B; 5-50 min, from 5% to 50% B; 50-55 min, from 50% to 98% B; 55-70 min, 98% B; 70-75 min, from 98% to 5% B; 75-90 min, 5% B at a flow rate of 14 ml min^-1^. The separation was monitored at 254 nm. NX-2 was collected at 43 min and the solvent evaporated in a vacuum concentrator (Christ, Osterode am Harz, Germany). NX-2 was further polished on Sephadex LH-20 (Sigma-Aldrich, Darmstadt, Germany) in a 460 × 15 mm column eluted isocratically with methanol at a flow rate of 1 ml min^-1^. The eluent was monitored at 254 nm.

### 5.5. Nuclear magnetic resonance (NMR) spectroscopy

1D (^1^H and ^13^C) NMR spectra were obtained from Bruker Avance 500 Hz and 2D (^1^H,^13^C-HSQC, ^1^H,^13^C-HMBC, ^1^H,^1^H-COSY) NMR spectra were obtained from an Agilent DD2 400 MHz system. The spectra were recorded at 500/400 MHz (^1^H) and 126/101 MHz (^13^C). Chemical shifts were referenced to internal TMS (dH 0, ^1^H) or MeOH-*d*4 (dc 49.0, ^13^C). The data are provided as **Supplementary Figures S1 to S5**.

## Supporting information

1H NMR spectrum of NX-2

13C NMR spectrum of NX-2

1H, 13C HSQC NMR spectrum of NX-2

1H, 13C HMBC spectrum of NX-2

1H, 1H COSY spectrum of NX-2

Comparison of NMR spectra of NX-2 with metabolite purified from F. culmorum 240.2sp

Conditions used for the amplification of TRI1 gene

Acc. nos. of sequences of the TRI1 gene

## Supplementary Data

Figure S1: ^1^H NMR spectrum of NX-2; Figure S2: ^13^C NMR spectrum of NX-2; Figure S3: ^1^H, ^13^C HSQC NMR spectrum of NX-2; Figure S4: ^1^H, ^13^C HMBC spectrum of NX-2; Figure S5: ^1^H, ^1^H COSY spectrum of NX-2. Table S1: Comparison of NMR spectra of NX-2 with metabolite purified from *F. culmorum* 240.2sp. Table S2: Conditions used for the amplification of *TRI1* gene. Table S3: Acc. nos. of sequences of the *TRI1* gene.

## Notes

### Competing Interest Statement

The authors have declared no competing interest.

