## Supplementary figures and images for "*Fusarium culmorum* produces NX-2 toxin simultaneously with deoxynivalenol and 3-acetyl-deoxynivalenol or nivalenol (submitted to Toxins)"

### 1H NMR spectrum of NX-2

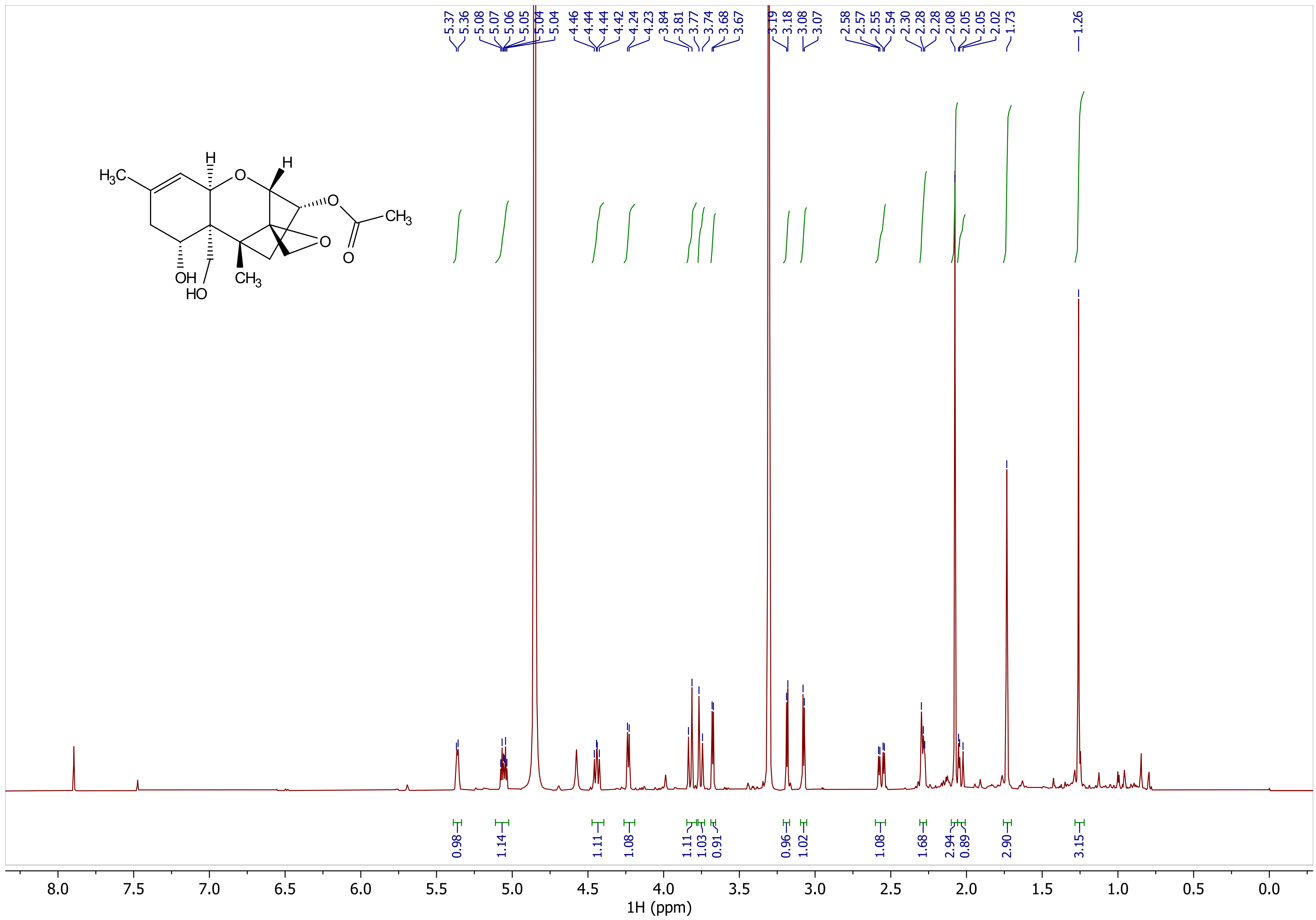

### 1H, 1H COSY spectrum of NX-2

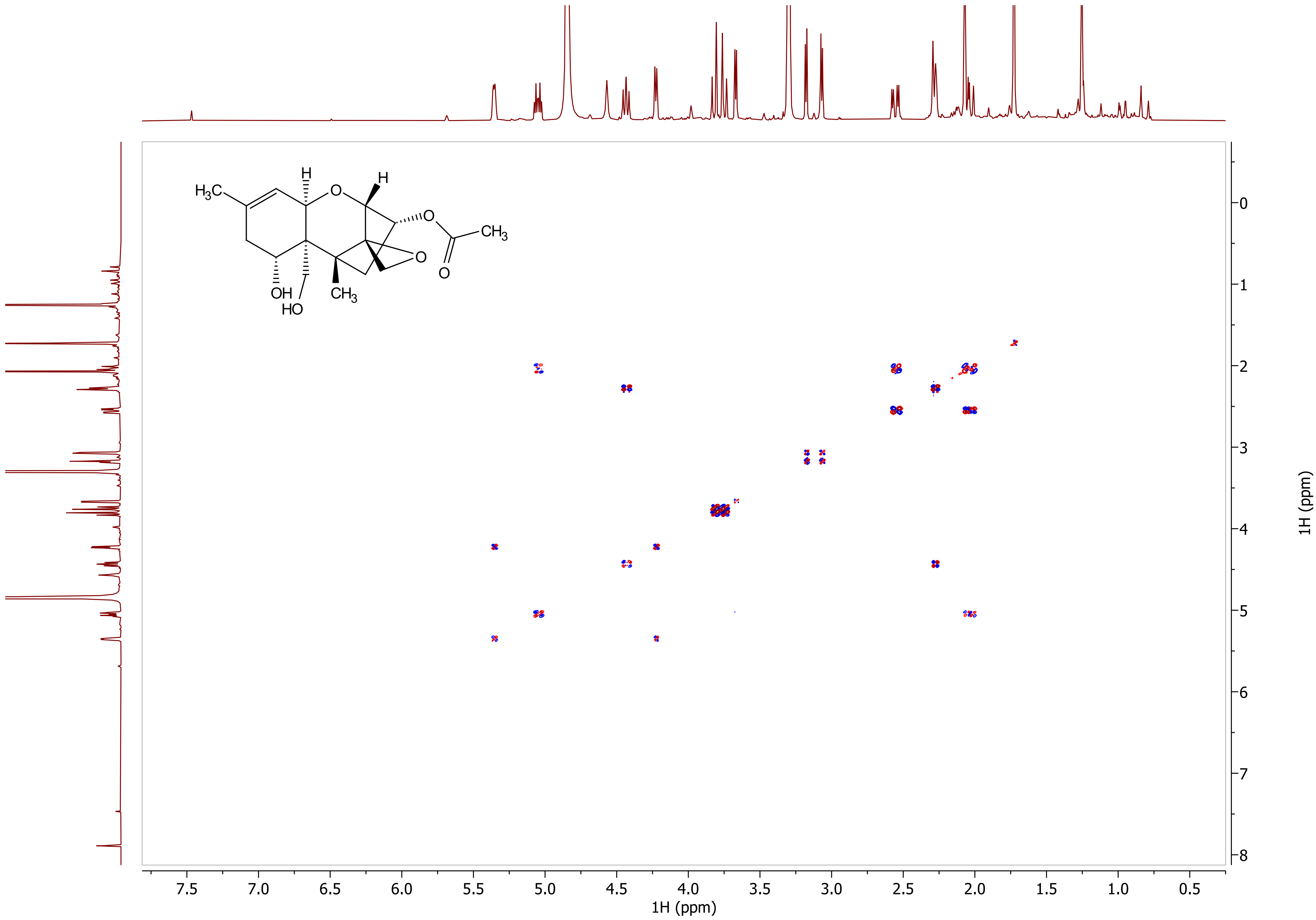

### 1H, 13C HMBC spectrum of NX-2

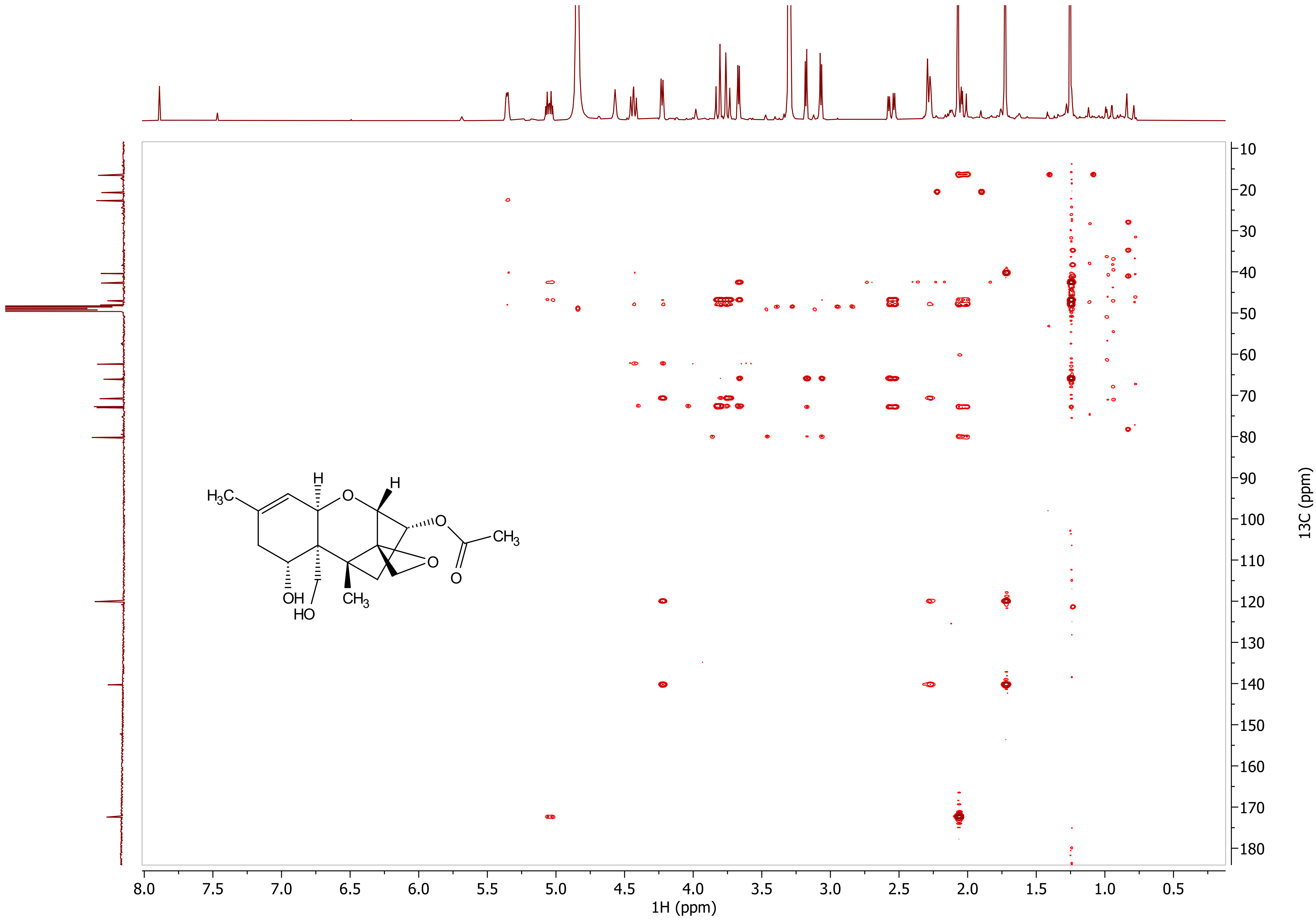

### 1H, 13C HSQC NMR spectrum of NX-2

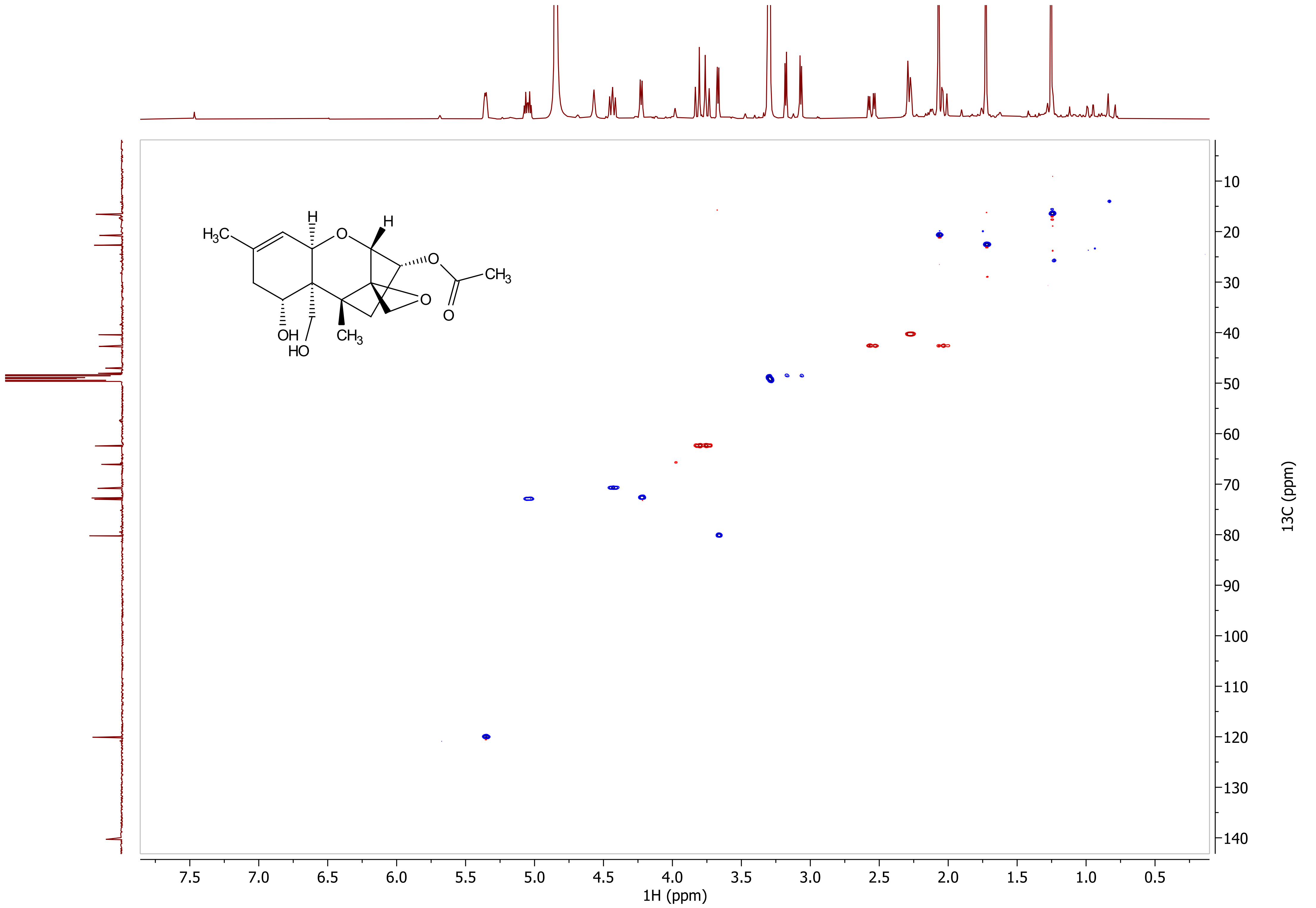

### 13C NMR spectrum of NX-2

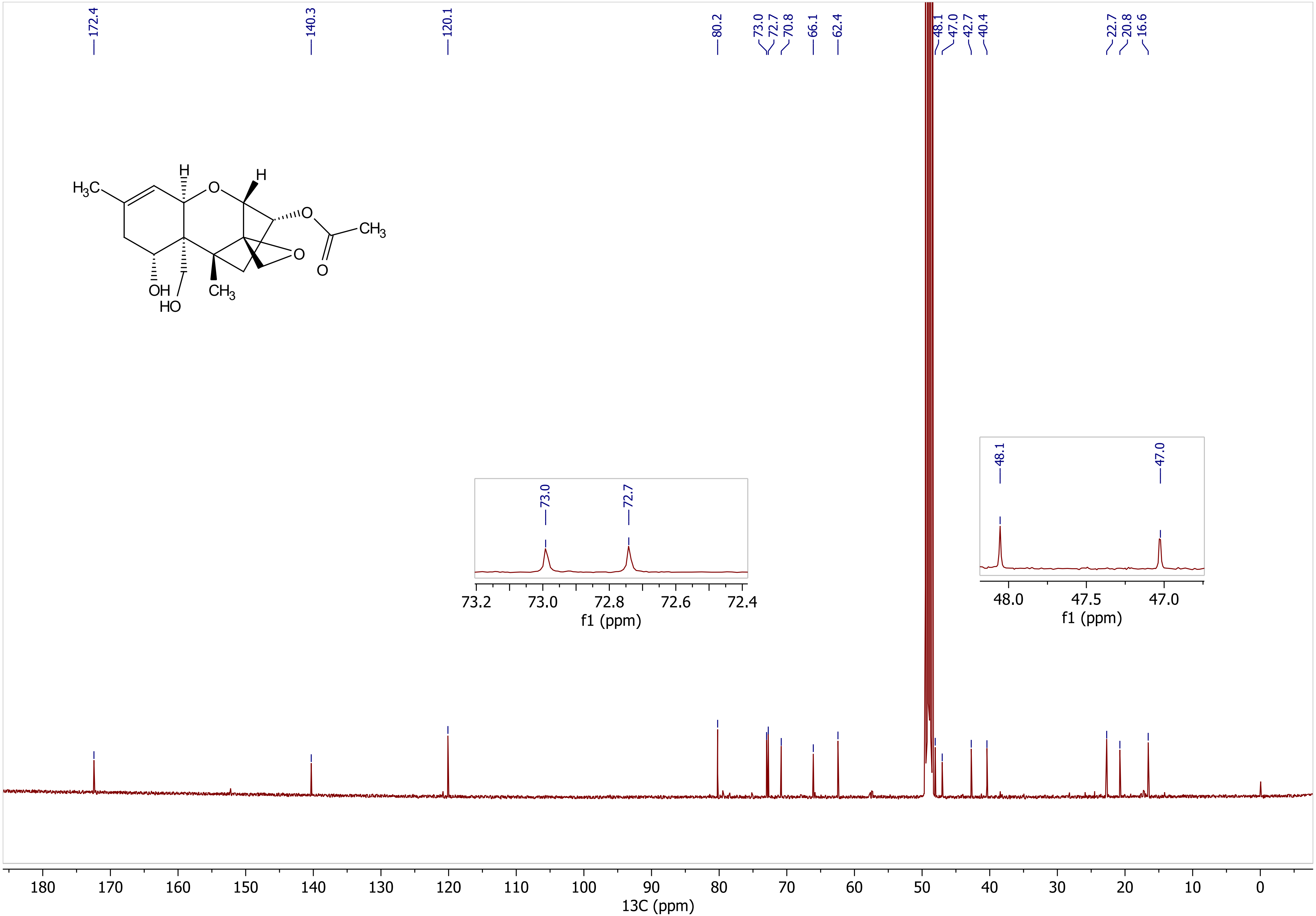
