## Supplementary material for "*Fusarium culmorum* produces NX-2 toxin simultaneously with deoxynivalenol and 3-acetyl-deoxynivalenol or nivalenol (submitted to Toxins)": Comparison of NMR spectra of NX-2 with metabolite purified from F. culmorum 240.2sp

**Table S1.** Comparison of published NMR spectral data for NX-2 with experimental data obtained on putative NX-2 from rice culture of *F. culmorum* 240.2.

| Position | Reported by Varga *et al.*, 2015 | | Experimental data | |
|---|---|---|---|---|
| | $\delta_C$ | $\delta_H$, mult. (J in Hz) | $\delta_C$ | $\delta_H$, mult. (J in Hz) |
| 2 | 80.3 | 3.68 d (4.4) | 80.2 | 3.68 d (4.4) |
| 3 | 73.1 | 5.06 dt (11.4, 4.4) | 73.0 | 5.06 dt (11.3, 4.4) |
| 4 | 42.9 | 2.56 dd (14.9, 4.4)<br>2.05[a] m | 42.7[b] | 2.56 dd (14.9, 4.4)<br>2.06 – 1.99 m |
| 5 | 47.2 | - | 47.0[b] | - |
| 6 | 48.2 | - | 48.1 | |
| 7 | 71.0 | 4.44 dd (9.0, 7.5) | 70.8[b] | 4.44 dd (9.0, 7.4) |
| 8 | 40.6 | 2.32–2.25 m | 40.4[b] | 2.32 – 2.25 m |
| 9 | 140.4 | | 140.3 | - |
| 10 | 120.2 | 5.36[a] m | 120.1 | 5.36 d (5.4) |
| 11 | 72.9 | 4.23 d (5.4) | 72.7[b] | 4.23 d (5.4) |
| 12 | 66.2 | - | 66.1 | - |
| 13 | 48.8 | 3.19 d (4.3)<br>3.08 d (4.3) | 48.6[b] | 3.18 d (4.4)<br>3.08 d (4.4) |
| 14 | 16.7 | 1.26 s | 16.6 | 1.26 s |
| 15 | 62.6 | 3.83 d (11.5)<br>3.76 d (11.5) | 62.4[b] | 3.82 d (11.6)<br>3.76 d (11.6) |
| 16 | 22.9 | 1.74 s | 22.7[b] | 1.73 s |
| 17 (3-Ac) | 172.6 | - | 172.4[b] | - |
| 18 (3-Ac) | 20.9 | 2.08 s | 20.8 | 2.08 s |

[a] misprint in [Varga et al.: Environ. Microbiol. 2015, 17:2588]; [b] all 13C peaks are shifted 0.1 ppm upfield and a 0.2 ppm difference comes from rounding the second decimal.
