## Supplementary material for "*Fusarium culmorum* produces NX-2 toxin simultaneously with deoxynivalenol and 3-acetyl-deoxynivalenol or nivalenol (submitted to Toxins)": Conditions used for the amplification of TRI1 gene

**Supplementary Table S2.** Conditions used for the amplification of *TRI1* gene of *F. culmorum*

| Forward primer | Reversed primer | Initial denaturation | Denaturation | Annealing | Elongation | Terminal elongation | No. of cycles | Length [bp] |
|---|---|---|---|---|---|---|---|---|
| TRI16IF1 | Tri1_5prime_RV | 95 °C, 30 s | 94 °C, 30 s | 59 °C, 30 s | 68 °C, 120 s | 68 °C, 5 min | 35 | 1641 |
| Tri1_5prime_FW | Tri1_SH_R | 95 °C, 30 s | 94 °C, 30 s | 59 °C, 30 s | 68 °C, 120 s | 68 °C, 5 min | 35 | 764 |
| Tri1_SH_F | FgTRI1-R1 | 95 °C, 30 s | 94 °C, 30 s | 59 °C, 30 s | 68 °C, 120 s | 68 °C, 5 min | 35 | 760 |
| FcTri1F | Tri1R | 95 °C, 30 s | 94 °C, 30 s | 58 °C, 30 s | 68 °C, 140 s | 68 °C, 5 min | 35 | 1745 |
