## Supplementary material for "*Fusarium culmorum* produces NX-2 toxin simultaneously with deoxynivalenol and 3-acetyl-deoxynivalenol or nivalenol (submitted to Toxins)": Acc. nos. of sequences of the TRI1 gene

**Table S2.** Accession numbers of sequences of the *TRI1* gene

| Species | Isolate | Accession No. |
|---|---|---|
| *F. culmorum* | K11.2 | OM144918 |
| *F. culmorum* | J31.2 | OM144919 |
| *F. culmorum* | IPP1000 | OM144920 |
| *F. culmorum* | IPP0999 | OM144921 |
| *F. culmorum* | IPP0619 | OM144922 |
| *F. culmorum* | IPP0618 | OM144923 |
| *F. culmorum* | IPP0213 | OM144924 |
| *F. culmorum* | IPP0212 | OM144925 |
| *F. culmorum* | IPP0211 | OM144926 |
| *F. culmorum* | DSM62188 | OM144927 |
| *F. culmorum* | 969 | OM144928 |
| *F. culmorum* | 966 | OM144929 |
| *F. culmorum* | 59.6st | OM144930 |
| *F. culmorum* | 55.6st | OM144931 |
| *F. culmorum* | 31.6st | OM144932 |
| *F. culmorum* | DSM62184 | OM144933 |
| *F. culmorum* | 3.37 | OM144934 |
| *F. culmorum* | 240.2sp | OM144935 |
| *F. culmorum* | 227. 2cst | OM144936 |
| *F. culmorum* | 215.1st | OM144937 |
| *F. graminearum* | CML3066 | LT222053 |
| *F. graminearum* | 06-267 | KX183401 |
| *F. graminearum* | 38383 | KX183278 |
| *F. graminearum* | 06-204 | KM999943 |
| *F. graminearum* | 02-264 | KM999941 |
| *F. graminearum* | 03-348 | KM999942 |
| *F. graminearum* | 40567 | KX183282 |
| *F. graminearum* | 45380 | KX183296 |
| *F. cerealis* (*F. crookwellense*) | 25805 | KX183232 |
| *F. pseudograminearum* | 28062 | KX183238 |
| *F. langsethiae* | NRRL53410 | HQ594538 |
| *F. sporotrichioides* | NRRL29977 | HQ594536 |
| *F. incarnatum* | NRRL31160 | GQ915526 |
| *F. sambucinum* | FRC R-07843 | GQ915521 |
| *F. poae* | FRC T-0962 | GQ915520 |
